# Hsp47 Promotes Biogenesis of Multi-subunit Neuroreceptors in the Endoplasmic Reticulum

**DOI:** 10.1101/2022.10.24.513629

**Authors:** Ya-Juan Wang, Xiao-Jing Di, Dong-Yun Han, Raad Nashmi, Brandon J. Henderson, Fraser J. Moss, Ting-Wei Mu

## Abstract

Protein homeostasis (proteostasis) deficiency is an important contributing factor to neurodegenerative, neurological, and metabolic diseases. However, how the proteostasis network orchestrates the folding and assembly of multi-subunit membrane proteins is not well understood. Previous proteomics studies identified Hsp47 (Gene: *SERPINH1*), a heat shock protein in the endoplasmic reticulum lumen, as the most enriched interacting chaperone for gamma-aminobutyric type A (GABA_A_) receptors. Here, we show that Hsp47 enhances neuronal GABA_A_ receptor functional surface expression, acting after Binding immunoglobulin Protein (BiP) to preferentially bind the folded conformation of GABA_A_ receptors. Therefore, Hsp47 promotes the subunit-subunit interaction, the receptor assembly process, and the anterograde trafficking of GABA_A_ receptors. These Hsp47 properties are also extended to other Cys-loop receptors, including nicotinic acetylcholine receptors. Therefore, in addition to its known function as a collagen chaperone, this work establishes that Hsp47 also plays a critical and general role in the maturation of multi-subunit neuroreceptors.

**Highlights:**

- Hsp47 positively regulates the functional surface expression of endogenous GABA_A_ receptors.
- Hsp47 acts after BiP and preferentially binds the folded conformation of GABA_A_ receptors.
- Hsp47 promotes the subunit-subunit assembly of GABA_A_ receptors.
- Hsp47 plays a critical and general role in the maturation of multi-subunit neuroreceptors.

## INTRODUCTION

Initially, the term “molecular chaperone” was coined to describe a nuclear protein that enables the assembly of nucleosomes from folded histone proteins and DNA [1, 2]. Since then, the role of chaperones, including the heat shock proteins, in facilitating protein folding [3, 4] and maintaining protein homeostasis (proteostasis) at the cellular, tissue, and organismal levels has been extensively explored [5–7]. Proteostasis deficiencies have been recognized in a growing number of neurodegenerative, neurological, and metabolic diseases [8–10]. Strategies to restore proteostasis, including applying regulators of the unfolded protein response (UPR) and Ca^2+^ regulation, have been actively developed to ameliorate such protein conformational diseases [11–15]. However, despite recent progress, the role of chaperones in regulating the folding and assembly of multi-subunit membrane proteins requires further elucidation [16, 17].

Multi-subunit membrane protein assembly in the endoplasmic reticulum (ER) is intimately linked to their folding and ER-associated degradation (ERAD). The current limited knowledge about the assembly process was gained from studying various classes of membrane proteins, including dimeric T cell receptors [18], trimeric P2X receptors [19], trimeric sodium channels [20], tetrameric potassium channels [21, 22], and pentameric nicotinic acetylcholine receptors (nAChRs) [23, 24]. We use γ-aminobutyric acid type A (GABA_A_) receptors as a physiologically important substrate to study their biogenesis [25]. GABA_A_ receptors are the primary inhibitory neurotransmitter-gated ion channels in mammalian central nervous systems [26] and provide most of the inhibitory tone to balance the tendency of excitatory neural circuits to induce hyperexcitability, thus maintaining the excitatory-inhibitory balance [27]. Functional GABA_A_ receptors are assembled as pentamers in the ER from eight subunit classes: α1-6, β1-3, γ1-3, δ, ε, θ, π, and ρ1-3. The most common subtype in the human brain contains two α1 subunits, two β2 subunits, and one γ2 subunit [28]. To form a heteropentamer, individual subunits need to fold into their native structures in the ER [29, 30] and assemble with other subunits correctly on the ER membrane [31, 32] (**Figure S1**). Only properly assembled pentameric receptors exit the ER, traffic through the Golgi for complex glycosylation, and reach the plasma membrane to perform their function. It was demonstrated that the α1 subunits fail to exit the ER on their own and are retained in the ER; after their assembly with β subunits, the α1β complex can exit the ER for subsequent trafficking to the plasma membrane [32, 33]. The inclusion of a γ2 subunit to form the pentamer further increases the conductance of the receptor and confers sensitivity to benzodiazepines [34]. Recently, it was reported that the synaptic localization of γ2-containing GABA_A_ receptors requires the LHFPL family protein LHFPL4 and Neuroligin-2 [35]. However, many of the fundamental questions about how the proteostasis network regulates the multi-subunit membrane protein assembly process remains to be determined.

Elucidating the proteostasis network for the subunit folding and assembly process of multi-subunit membrane proteins and their biogenesis pathway in general, is important to fine-tune their function in physiological and pathological conditions. Loss of function of GABA_A_ receptors is one prominent cause of genetic epilepsies [36, 37]. Furthermore, numerous variations in a single subunit cause subunit protein misfolding in the ER and/or disrupt assembly of the pentameric complex, leading to excessive ERAD, decrease cell surface localization of the receptor complex, and result in imbalanced neural circuits [25, 38]. The elucidation of the GABA_A_ receptor proteostasis network will guide future efforts to develop strategies that restore proteostasis of variant GABA_A_ receptors to ameliorate corresponding diseases, such as genetic epilepsies.

Recently, our quantitative affinity purification mass spectrometry-based proteomics analysis identified Hsp47 (Gene: *SERPINH1*) as the most enriched GABA_A_ receptor-interacting chaperone [39]. Hsp47 is an ER-resident protein with a RDEL (Arg-Asp-Glu-Leu) ER retention signal [40, 41]. Among the large Serpin (*ser*ine *p*rotease *in*hibitor) superfamily, Hsp47 is the only one reported to show a molecular chaperone function [42]. Current literature describes Hsp47 as a collagen-specific chaperone [43, 44]. However, its broader role has been indicated [45], such as interacting with the inositol-requiring enzyme 1α (IRE1α) to regulate the UPR [46] as well as interacting with amyloid precursor protein (APP) in the central nervous system (CNS) [47]. Here, we demonstrate that Hsp47 enhances the functional surface expression of both endogenous GABA_A_ receptors, and other Cys-loop receptors in the CNS. Furthermore, a mechanistic study revealed that Hsp47 promotes the folding and assembly of multi-subunit neuroreceptors in the ER. Consequently, our results support a general role of Hsp47 in multi-subunit membrane protein quality control.

## RESULTS

### Hsp47 directly interacts with GABA_A_ receptor subunits

Since we previously identified Hsp47 as the most enriched GABA_A_ receptor-interacting chaperone in HEK293T cells using quantitative proteomics [39], here we evaluated the interaction between Hsp47 and GABA_A_ receptors in more detail. Co-immunoprecipitation assays using mouse brain homogenates showed that the endogenous Hsp47 binds to endogenous GABA_A_ receptor α1 subunits in the CNS (**Figure 1A**). Furthermore, to test a direct interaction between Hsp47 and GABA_A_ receptor subunits, we carried out an *in vitro* binding assay using recombinant GST-tagged α1 or β2 subunits and recombinant His-tagged Hsp47. The anti-His antibody pulldown detected the α1 subunit in the GST-α1 complex (**Figure 1B**, lane 5) and the β2 subunit in the GST-β2 complex (**Figure 1B**, lane 10). No α1 or β2 bands were detected in the GST control complex (**Figure 1B**, lanes 4 and 9), indicating that Hsp47 directly binds to the GABA_A_ receptor α1 and β2 subunits *in vitro*.

**Figure 1.**
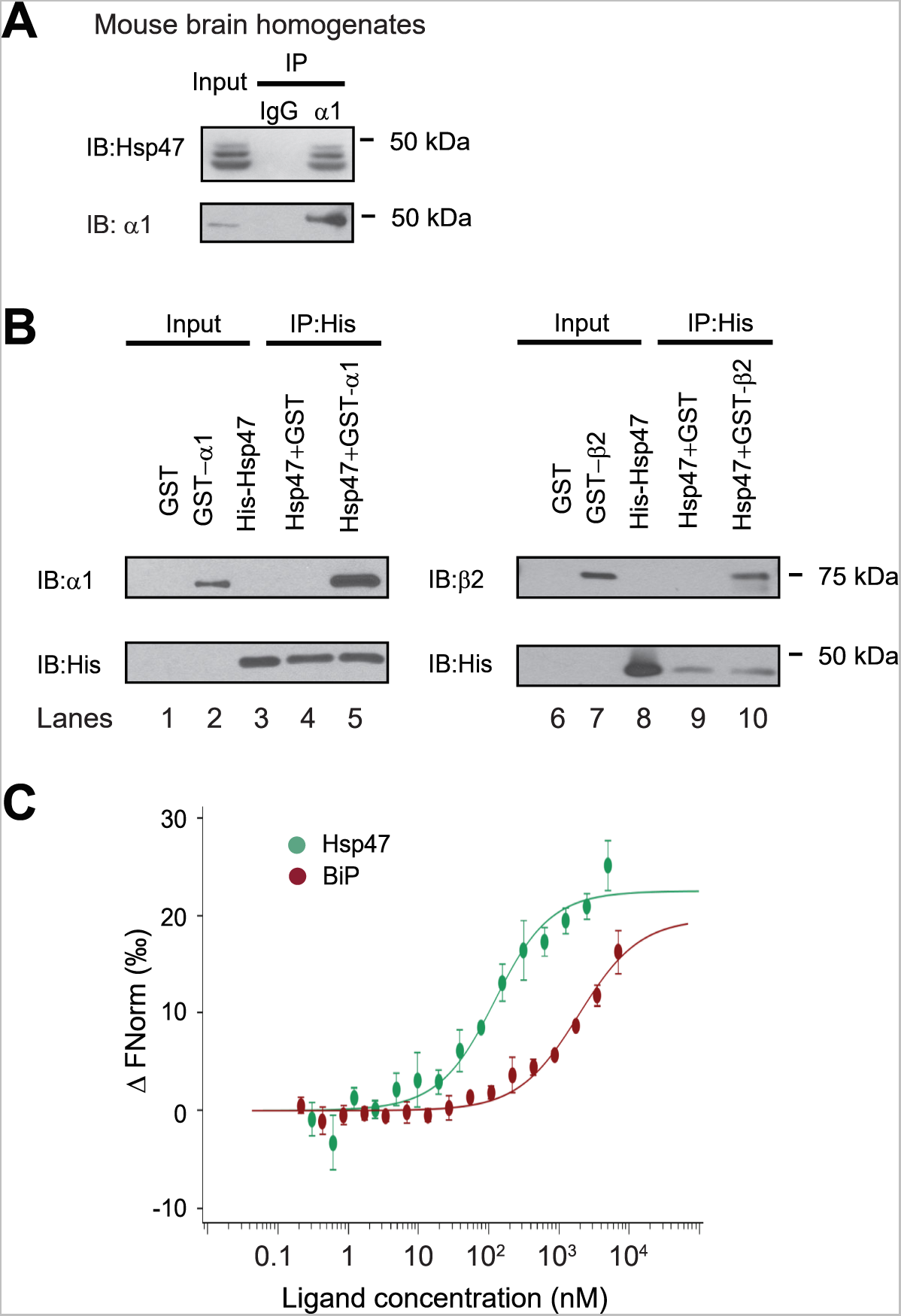
Hsp47 interacts with GABA_A_ receptors. (**A**) Endogenous interactions between GABA_A_ receptor α1 subunits and Hsp47. Mouse brain homogenates from 8-10 weeks C57BL/6J mice were immunoprecipitated with an anti-α1 antibody, and the immunoisolated eluents were blotted with indicated antibodies (n = 3). IgG was included as a negative control for non-specific binding. (**B**) Recombinant Hsp47 binds recombinant α1 subunit and β2 subunit of GABA_A_ receptors *in vitro*. GST, GST-tagged α1 or GST-tagged β2 recombinant protein was mixed with His-tagged Hsp47 in buffers containing 1% Triton X-100. The protein complex was isolated by immunoprecipitation using an anti-His antibody, and the immunopurified eluents were separated by SDS-PAGE and blotted with indicated antibodies (n = 3). (**C**) MicroScale Thermophoresis (MST) was used to determine the binding affinities between ER luminal chaperones (Hsp47 and BiP) to RED-labeled His-α1(ERD). Increasing concentrations of recombinant Hsp47 or BiP proteins (0.20 nM – 10 μM) were incubated with 50 nM RED-labeled His-α1(ERD) protein in PBS with Tween-20 (0.05%) (n = 3). Then samples were loaded to the capillaries and measured using a Monolith NT.115 instrument with the settings of 40% LED/excitation and 40% MST power. The data were analyzed using the Monolith software for the calculation of the dissociation constant (Kd). IP, immunoprecipitation; IB, immunoblotting.

Since Hsp47 resides in the ER lumen, it presumably interacts with the GABA_A_ receptor ER luminal domain (ERD). According to a circular dichroism study, the α1 subunit ERD domain adopts a well-defined secondary structure [48]. Therefore, we determined the binding affinity between α1(ERD) and Hsp47. A MicroScale Thermophoresis (MST) assay reported a strong interaction (dissociation constant, Kd = 102 `±` 10 nM) between α1(ERD) and Hsp47 (**Figure 1C**). In addition, due to the established role of Binding immunoglobulin Protein (BiP), a Hsp70 family chaperone in the ER lumen, in binding GABA_A_ receptors [49], we measured the binding affinity between α1(ERD) and BiP, showing a Kd = 1906 ± 210 nM (**Figure 1C**). Therefore, Hsp47 binds more strongly to GABA_A_ receptors compared to BiP, possibly because Hsp47 and BiP bind different GABA_A_ receptor folding states (see below).

### Hsp47 positively regulates the functional surface expression of endogenous GABA_A_ receptors in neurons

To the best of our knowledge, functional regulation of GABA_A_ receptor and other ion channels by Hsp47 in the CNS has not been previously reported. Hsp47 is widely distributed in the CNS, including the cortex, hippocampus, hypothalamus, cerebellum, and olfactory bulb tested (**Figure S2A**), which is consistent with the report that Hsp47 is robustly detected in primary cortical and hippocampal neurons and brain slices [47]. Concomitantly, GABA_A_ receptors are also distributed in these brain areas (**Figure S2A**) [28, 50].

Because GABA_A_ receptors must reach the plasma membrane to act as ligand-gated ion channels, we first performed an indirect immunofluorescence microscopy experiment to evaluate how Hsp47 regulates their endogenous surface expression levels in primary rat hippocampal neurons. The application of anti-GABA_A_ receptor subunit antibodies that recognize their extracellular N-termini without a prior membrane permeabilization step enabled us to label only the cell surface expressed proteins. Transduction of lentivirus carrying Hsp47 siRNA led to substantial depletion of Hsp47 in neurons (**Figure S2B**), and knocking down Hsp47 significantly decreased the surface staining of the major subunits of GABA_A_ receptors, including the α1 subunits and β2/β3 subunits (**Figure 2A**, cf. column 2 to column 1). This result indicated that Hsp47 positively regulates the surface protein levels of endogenous GABA_A_ receptors. Furthermore, whole-cell patch-clamp electrophysiology recordings demonstrated that depleting Hsp47 significantly decreased the peak GABA-induced currents from 1660 ± 413 pA in the presence of scrambled siRNA to 886 ± 157 pA after the application of lentivirus carrying Hsp47 siRNA in hippocampal neurons (**Figure 2B**). Collectively, the experiments in **Figure 2** unambiguously reveal a novel role of Hsp47 as a positive regulator of the functional surface expression of endogenous GABA_A_ receptors, an important neuroreceptor.

**Figure 2.**
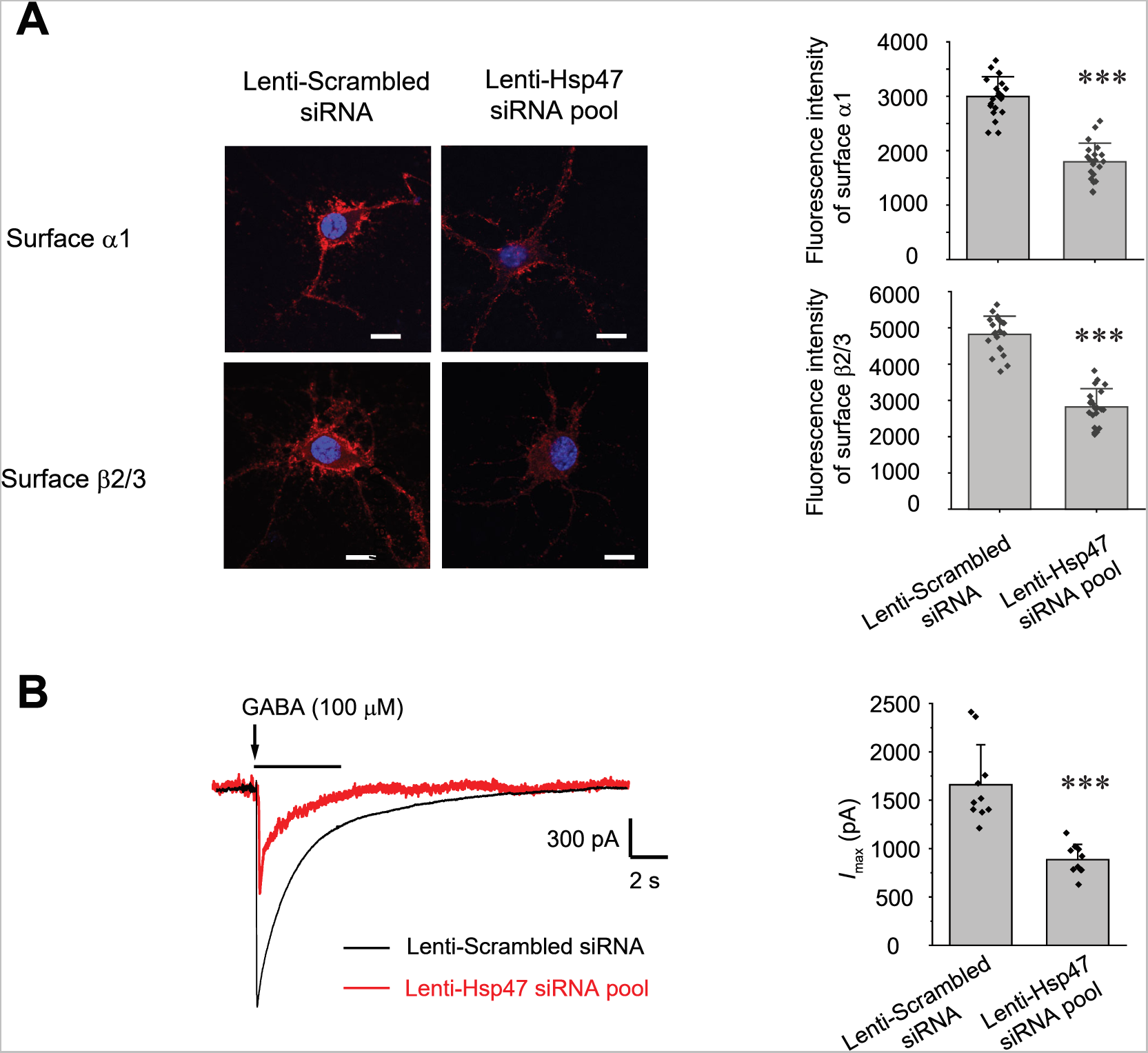
Hsp47 positively regulates the surface expression of endogenous GABA_A_ receptors in cultured neurons. (**A**) Effect of Hsp47 on the surface expression of endogenous GABA_A_ receptor subunits in primary rat hippocampal neurons. Cultured neurons were transduced with Hsp47 siRNA lentivirus or scrambled siRNA lentivirus at days *in vitro* (DIV) 10. Forty-eight hours post transduction, surface GABA_A_ receptors were stained using anti-α1 subunit or anti-β2/β3 subunit antibodies without membrane permeabilization. The cells were then washed, and permeabilized before we stained the nuclei with DAPI. At least 20 neurons from at least three transductions were imaged by confocal microscopy for each condition. Representative images are shown on the left side of panel A. Scale bar = 10 μm. Quantification of the fluorescence intensity of the surface GABA_A_ receptor subunits after background correction was shown on the right. (**B**) Whole-cell patch clamping was performed to record GABA-induced currents. Neurons were subjected to transduction as in (**A**). The recordings were carried out 48 hours post transduction. Representative traces are shown in the left-hand panel. Peak current amplitude (*I*_max_) is shown on the right (n = 10). The holding potential was set at −60 mV. pA: picoampere. Each data point is reported as mean ± SD. Statistical significance was calculated using an unpaired two-tailed Student’s t-Test. *** *p* < 0.001.

### Hsp47 preferentially binds the folded conformation of GABA_A_ receptor subunits and promotes their ER-to-Golgi trafficking

Since Hsp47 is an ER luminal chaperone, we hypothesized that to enhance the surface trafficking of GABA_A_ receptors, Hsp47 promotes their protein folding in the ER and subsequent anterograde trafficking. We used an endoglycosidase H (Endo H) enzyme digestion assay to monitor the ER-to-Golgi trafficking of GABA_A_ receptors, also as a surrogate to determine whether GABA_A_ receptors are folded and assembled properly in the ER [49]. The Endo H enzyme selectively cleaves after asparaginyl-*N*-acetyl-D-glucosamine (GlcNAc) in the N-linked glycans in the ER, but it cannot remove this oligosaccharide chain after the high mannose form is enzymatically remodeled in the Golgi. Therefore, Endo H resistant subunit bands represent properly folded and assembled, post-ER subunit glycoforms, which traffic at least to the Golgi. Due to the heterogeneity of GABA_A_ receptor subunits in neurons, we employed HEK293T cells to exogenously express the major subtype of GABA_A_ receptors containing α1, β2, and γ2 subunits for the mechanistic study [51]. The Endo H digestion assay showed that Hsp47 overexpression increased the Endo H resistant band intensity (**Figure 3A**, cf. lane 4 to lane 2) and the ER to Golgi trafficking efficiency, represented by the ratio of the Endo H resistant α1 band / total α1 band (**Figure 3A**, cf. lane 4 to lane 2; quantification shown in the **Figure 3A** bottom panel). This result indicated that Hsp47 enhanced the folding and assembly of GABA_A_ receptors in the ER and thus their ER-to-Golgi trafficking.

**Figure 3.**
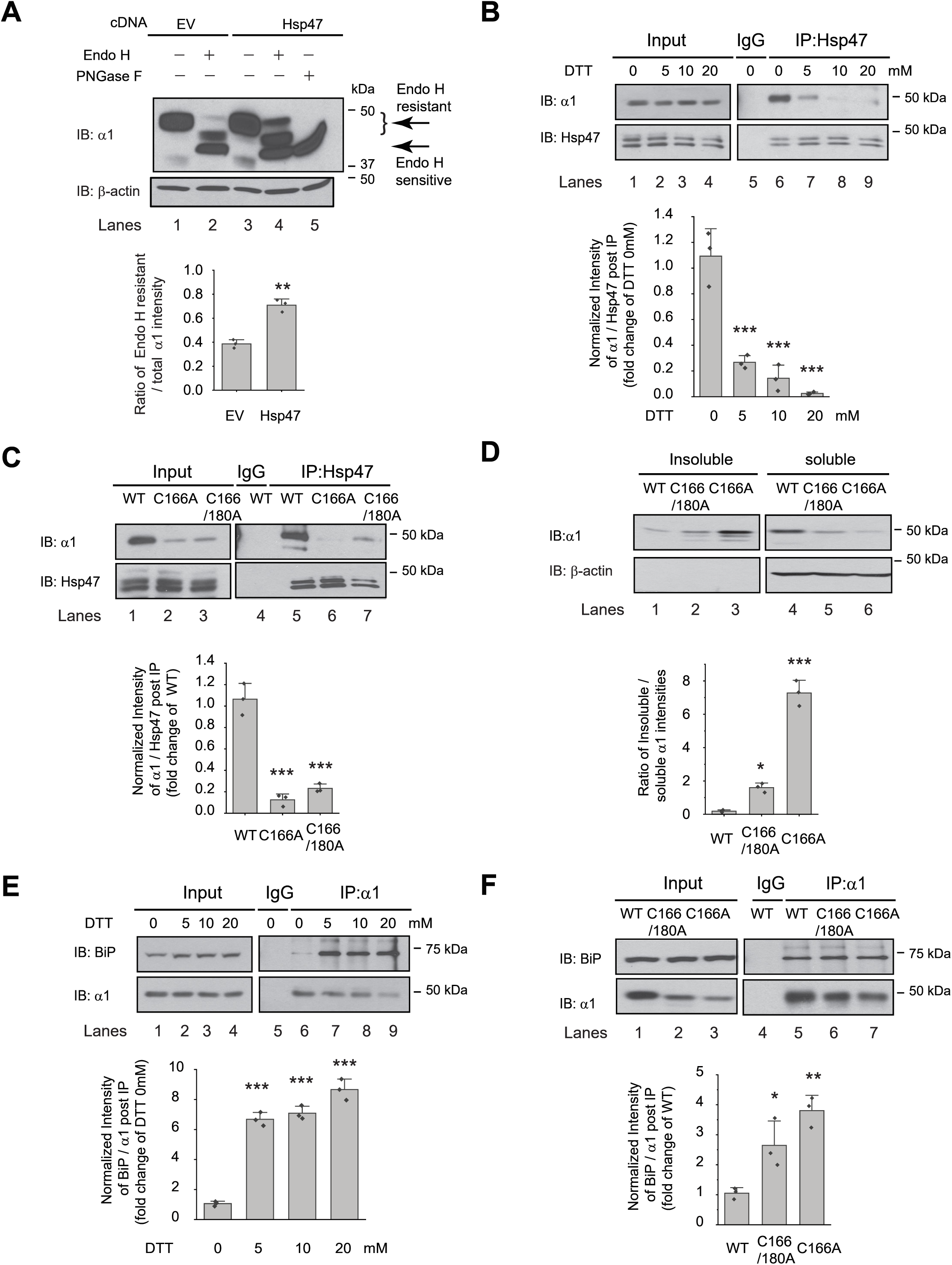
Hsp47 preferentially binds the folded conformation of GABA_A_ receptor subunits. (**A**) Overexpression of Hsp47 increases the endo H-resistant post-ER glycoform of the α1 subunit in HEK293T cells stably expressing α1β2γ2 GABA_A_ receptors. PNGase F treatment serves as a control for unglycosylated α1 subunit (lane 5). Two endo H-resistant bands were detected for the α1 subunit, indicated by the bracket (lanes 2 and 4). Quantification of the ratio of endo H- resistant / total α1 subunit bands, as a measure of the ER-to-Golgi trafficking efficiency, is shown on the bottom (n = 3). (**B**) Dithiothreitol (DTT) treatment decreases the interaction between Hsp47 and α1 subunit of GABA_A_ receptors. HEK293T cells stably expressing WT α1β2γ2 GABA_A_ receptors were treated with indicated concentration of DTT in the PBS buffer for 10 minutes. Then Triton X-100 cell extracts were immunoprecipitated with a mouse anti- Hsp47 antibody, and the immunoisolated eluents were subjected for immunoblotting assay (n = 3). Quantification of the relative intensity of α1/Hsp47 post IP, as a measure of their interactions, is shown on the bottom panel. (**C**) Disulfide bond mutations in the α1 subunit decrease the interaction between Hsp47 and α1 subunit of GABA_A_ receptors. HEK293T cells were transiently transfected with WT α1β2γ2, α1(C166A)β2γ2, or α1(C166A, C180A)β2γ2 subunits. Forty-eight hours post transfection, Triton X-100 cell extracts were immunoprecipitated with a mouse anti- Hsp47 antibody, and the immunoisolated eluents were subjected for immunoblotting assay (n = 3). Quantification of the relative intensity of α1/Hsp47 post IP is shown on the bottom panel. (**D**) Disulfide bond mutations in the α1 subunits decrease the solubility of the α1 subunit protein. HEK293T cells were transiently transfected as in (**C**). Forty-eight hours post transfection, the Triton X-100 detergent soluble fractions and the Triton X-100 detergent insoluble fractions were isolated for immunoblotting assay (n = 3). Quantification of the ratio of insoluble/soluble fractions, as a measure of relative aggregation, is shown on the bottom panel. (**E**) DTT treatment increases the interaction between BiP and α1 subunit of GABA_A_ receptors. HEK293T cells stably expressing α1β2γ2 GABA_A_ receptors were treated with indicated concentrations of DTT in PBS for 10 minutes. Then Triton X-100 cell extracts were immunoprecipitated with a mouse anti-α1 antibody, and the immunoisolated eluents were subjected for immunoblotting assay (n = 3). Quantification of the relative intensity of BiP/α1 post IP is shown on the bottom panel. (**F**) The disulfide mutations of α1 subunit increase the interaction between BiP and the α1 subunit. HEK293T cells were transiently transfected as in (**C**). Forty-eight hours post transfection, Triton X-100 cell extracts were immunoprecipitated with a mouse anti-α1 antibody, and the immunoisolated eluents were subjected for immunoblotting assay (n = 3). Quantification of the relative intensity of BiP/α1 post IP is shown on the bottom panel. IP, immunoprecipitation; IB, immunoblotting. Each data point is reported as mean ± SD. Significant difference was analyzed by t-test (**A**), or a one-way ANOVA followed by post hoc Tukey’s HSD test (**B-F**). *, *p* < 0.05; **, *p* < 0.01; ***, *p* < 0.001.

We next evaluated how Hsp47 coordinates the folding and assembly of GABA_A_ receptors. We perturbed α1 subunit folding both genetically and chemically in HEK293T cells expressing GABA_A_ receptors. We then evaluated the correlation between the relative folding degree of the α1 subunits and their interaction with Hsp47 using a co-immunoprecipitation assay. Individual GABA_A_ receptor subunits have a signature disulfide bond in their large N-terminal domain. Adding dithiothreitol (DTT), which is cell-permeable, to the cell culture media for 10 min destroyed the signature disulfide bond between Cys166 and Cys180 in the α1 subunit, thus compromising its folding. This operation did not change the total α1 protein levels (**Figure 3B**, cf. lanes 2-4 to lane 1) possibly because, during such a short time, degradation of the misfolded α1 subunit was not substantial. In sharp contrast, adding DTT significantly decreased the α1 protein that was pulled down by Hsp47 in a dose-dependent manner (**Figure 3B,** cf. lanes 7-9 to lane 6, quantification shown in the bottom panel). This indicates that eliminating the signature disulfide bond in the α1 subunit decreased its interaction with Hsp47 and supports the hypothesis that Hsp47 preferentially binds to the folded α1 subunit conformation.

In addition, we genetically disrupted the signature disulfide bond in the α1 subunit either by introducing a single C166A mutation or C166A/C180A double mutations. The co-immunoprecipitation assay clearly demonstrated that both the single and double mutations led to a decreased interaction between the α1 subunit and Hsp47 (**Figure 3C**, cf. lanes 6 and 7 to lane 5, quantification shown in the bottom panel). To evaluate the relative conformational stability of the α1 subunit variants, we examined the Triton X-100 detergent soluble fractions and the Triton X-100 detergent-insoluble fractions. The percentage of the insoluble fractions in the C166A single mutation and the C166A/C180A double mutations is significantly greater than that in the WT receptors (**Figure 3D**, quantification shown in the bottom panel), indicating that disrupting the signature disulfide bond induces aggregation. Notably, the C166A single mutant is more prone to aggregation than the C166A/C180A double mutant, which is not surprising because the single C166A mutant subunit retains an unpaired Cys180 in the ER lumen that remains available for cross-linking.

During the biogenesis in the ER, GABA_A_ receptors need to interact with a network of chaperones and folding enzymes, such as BiP, to acquire their native structures [39]. BiP, which binds the hydrophobic patches of unfolded proteins and prevents their aggregation [52, 53], interacts with the GABA_A_ receptors in the ER [32, 49]. We reasoned that if Hsp47 preferentially binds the folded conformation of the GABA_A_ receptor subunits, it would act after BiP because BiP is expected to act early in the protein folding step in the ER [54, 55]. We therefore evaluated how disrupting appropriate α1 subunit folding influenced their interaction with BiP. As hypothesized, the interactions between the α1 subunits and BiP were significantly enhanced when the signature disulfide bonds were chemically destroyed by adding DTT to the cell culture media (**Figure 3E,** cf. lanes 7, 8, and 9 to lane 6, quantification shown in the bottom panel). Genetic destruction of the α1 subunit disulfide bonds in the C166A single mutant or the C166A/C180A double mutant (**Figure 3F** cf. lanes 6 and 7 to lane 5, quantification shown in the bottom panel) produced similar results. Collectively, **Figure 3** indicates that inducing misfolding of GABA_A_ receptors compromises their interactions with Hsp47, whereas, in sharp contrast, enhances their interactions with BiP. Therefore, BiP preferentially binds the unfolded/misfolded states, whereas Hsp47 preferentially binds the properly folded states of the α1 subunits. Hsp47 must therefore acts after BiP to enhance the productive folding of GABA_A_ receptors.

### Hsp47 enhances the subunit-subunit assembly of GABA_A_ receptors

A cellular environment is required for the assembly of the majority of ion channels [23], indicating that factors other than ion channel subunits themselves are necessary in this process. We next tested our hypothesis that Hsp47 promotes efficient GABA_A_ receptor subunit assembly. We used Förster resonance energy transfer (FRET) to evaluate the cellular interactions between GABA_A_ receptor subunits. We incorporated enhanced cyan fluorescent protein (CFP) (donor) into the TM3-TM4 intracellular loop of the α1 subunit and enhanced yellow fluorescent protein (YFP) (acceptor) into the TM3-TM4 intracellular loop of the β2 subunit. The addition of CFP/YFP into the large intracellular loops of GABA_A_ receptors did not change the function of GABA_A_ receptors since dose response to GABA was indistinguishable between α1β2γ2 receptors and (CFP-α1)(YFP-β2)γ2 receptors according to patch-clamp electrophysiology recordings in HEK293T cells (**Figure S3**). The TM3-TM4 intracellular loops are the most variable segment within GABA_A_ receptor subunits and their splice variants. These were often replaced with short sequences in structural studies [56, 57]. Intracellular loops that incorporated CFP or YFP were also utilized in FRET experiments performed on nAChRs [58], members of the same Cys-loop superfamily to which GABA_A_ receptors belong. Pixel-based FRET experiments showed that mean FRET efficiency was 24.4 ± 3.1% for (CFP-α1)(YFP-β2)γ2 receptors (**Figure 4A**, row 1, column 3); overexpressing Hsp47 significantly increased the mean FRET efficiency to 31.7 ± 5.7% (**Figure 4A**, row 2, column 3), indicating that Hsp47 positively regulates the assembly between α1 and β2 subunits of GABA_A_ receptors. In addition, the co-immunoprecipitation assay showed that overexpression of Hsp47 significantly increased the relative amount of the β2 subunit that was pulled down with the α1 subunit (**Figure 4B**, cf. lane 5 to 4, quantification shown on the bottom), indicating that Hsp47 promotes incorporation of β2 subunits into GABA_A_ receptor pentamers.

**Figure 4.**
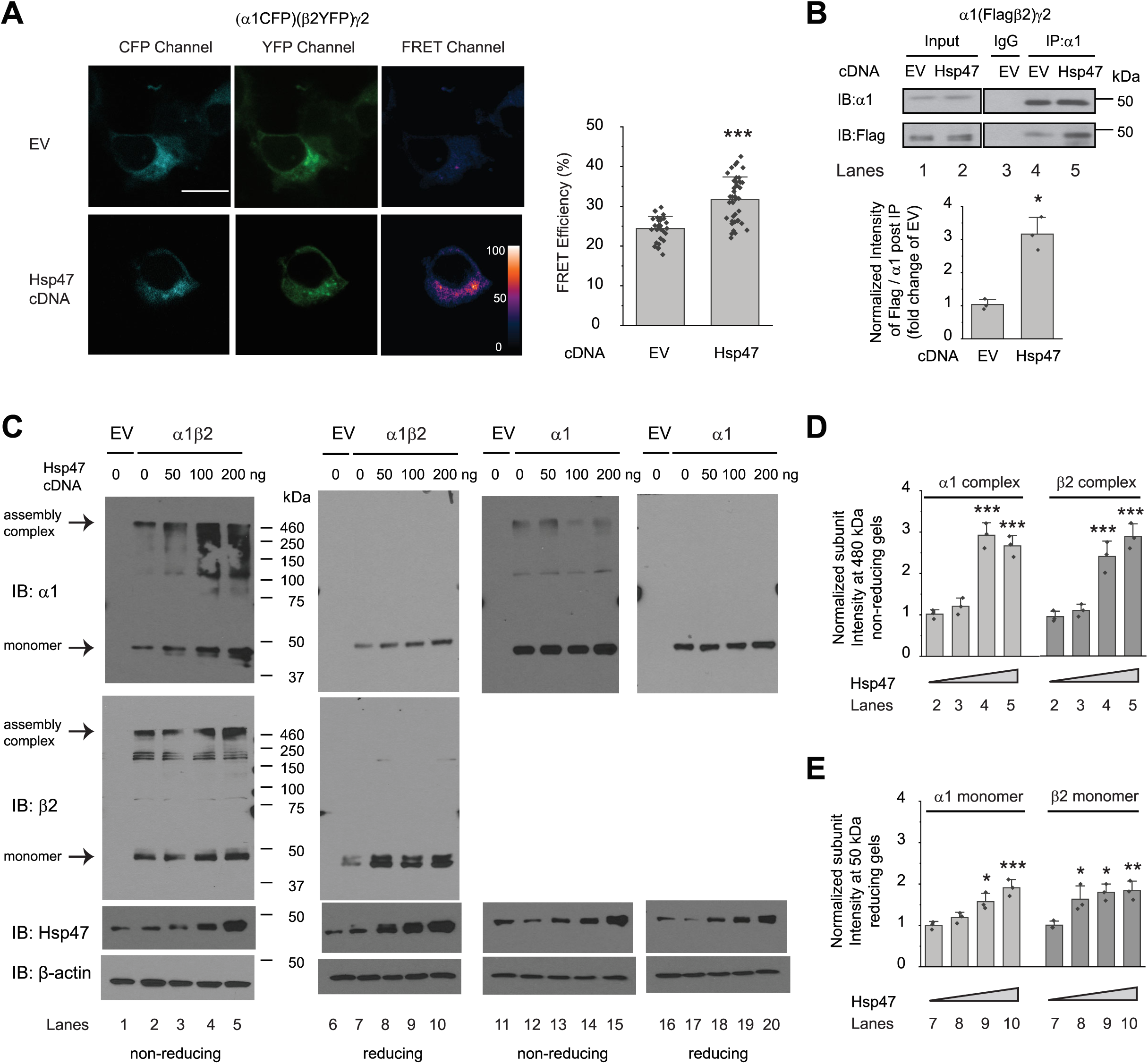
Hsp47 promotes the assembly of GABA_A_ receptors. (**A**) Hsp47 overexpression increases FRET efficiency between CFP-tagged α1 subunit and YFP-tagged β2 subunit of GABA_A_ receptors. HEK293T cells were transfected with CFP-tagged α1 subunit, YFP-tagged β2 subunit, and γ2 subunit; in addition, cells were transfected with empty vector (EV) control or Hsp47 cDNA. Forty-eight hours post transfection, pixel-based FRET was used to measure the FRET efficiency between α1-CFP and β2-YFP by using a confocal microscope. Representative images were shown for the CFP channel (1st columns), YFP channel (2nd columns), and FRET efficiency (3rd columns). Scale bar = 10 μm. Quantification of the FRET efficiency from 30-41 cells was achieved using the ImageJ PixFRET plug-in, and shown on the right. (**B**) Overexpression of Hsp47 increases the interaction between α1 and β2 subunit of GABA_A_ receptors. HEK293T cells stably expressing α1(Flag-β2)γ2 GABA_A_ receptors were transfected with empty vector (EV) control or Hsp47 cDNA. Forty-eight hours post transfection, Triton X-100 cell extracts were immunoprecipitated with a mouse anti-α1 antibody, and the immunoisolated eluents were subjected to immunoblotting assay. Quantification of the relative intensity of Flag-β2 / α1 post IP is shown on the bottom (n = 3). (**C**) HEK293T cells were transiently transfected with empty vector (EV), α1 subunits alone, or both α1 and β2 subunits of GABA_A_ receptors together with Hsp47 cDNA plasmids at various concentrations. Forty-eight hours post transfection, cells were lysed in RIPA buffer, and the total cell lysates were subjected to SDS-PAGE under non-reducing conditions and reducing conditions and immunoblotting analysis. (**D**) Quantification of the 480 kDa band intensities for α1 and β2 subunits under non- reducing conditions (lanes 2-5 in **C**) (*n* = 3). (**E**) Quantification of the 50 kDa band intensities for α1 and β2 subunits under reducing conditions (lanes 7-10 in **C**) (*n* = 3). IP, immunoprecipitation; IB, immunoblotting. Each data point is reported as mean ± SD. Significant difference was analyzed by t-test (**A**, **B**) or a one-way ANOVA followed by post hoc Tukey’s HSD test (**D**, **E**). *, *p* < 0.05; **, *p* < 0.01; ***, *p* < 0.001.

Further, we used non-reducing protein gels to evaluate how Hsp47 influences the formation of the oligomeric subunits during the assembly process in the ER. The absence of reducing reagents in the protein gel’s sample loading buffer preserves the intra- and inter-subunit disulfide bonds, which is expected to enable the detection of subunit oligomerization. In HEK293T cells expressing α1β2 receptors, distinct bands around 480 kDa were visible for both α1 subunit and β2 subunit in non-reducing gels (**Figure 4C**, lanes 2 to 5), indicating that the 480 kDa complex corresponds to the α1β2 hetero-oligomers. Moreover, the apparent molecular weight of the 480 kDa complex agrees with the molecular weight of the detected native GABA_A_ receptors obtained from the cerebellum using blue native protein gels [35]. Therefore, the 480 kDa complex probably corresponds to the correctly assembled receptor complex. Strikingly, overexpression of Hsp47 increased the intensity of the 480 kDa bands for both α1 and β2 subunits (**Figure 4C**, cf. lanes 3-5 to lane 2, quantification in **Figure 4D**), indicating that Hsp47 promotes the formation of the properly assembled oligomers. In addition, reducing protein gels showed that overexpressing Hsp47 increased the band intensities for α1 and β2 subunits at 50 kDa in HEK293T cells expressing α1β2 receptors (**Figure 4C**, cf. lanes 8 to 10 to lane 7, quantification in **Figure 4E**). Moreover, we carried out control experiments using HEK293T cells expressing only α1 subunits because α1 subunits alone cannot exit the ER [33]. Non-reducing gels revealed that the majority of the detected α1 protein was in the monomeric form (**Figure 4C**, lane 12), and overexpression of Hsp47 did not change the intensity of α1 subunit band on the non-reducing gels (**Figure 4C**, cf. lanes 13-15 to lane 12) or using reducing gels (**Figure 4C**, cf. lanes 18-20 to lane 17). This probably occurred because α1 subunits alone cannot assemble to form a trafficking-competent complex to exit the ER. Collectively, these results indicated that Hsp47 promotes the assembly of the native pentameric GABA_A_ receptor complexes for their subsequent ER exit and trafficking to the Golgi and plasma membrane.

### Hsp47 rescues the function of epilepsy-associated GABA_A_ receptors carrying the α1(A322D) variant

We next evaluated the effect of Hsp47 on the trafficking and function of pathogenic GABA_A_ receptors using a well-characterized misfolding-prone α1(A322D) variant [59]. The A322D mutation introduces an extra negative charge in the TM3 helix of the α1 subunit, leading to inefficient insertion of TM3 into the lipid bilayer and fast degradation of the α1 subunit [49, 59]. As a result, the α1(A322D) variant causes loss of function of GABA_A_ receptors and juvenile myoclonic epilepsy [60]. An Endo H enzyme digestion assay showed that overexpressing Hsp47 increased the ratio of Endo H resistant α1 / total α1 bands from 0.20 ± 0.03 to 0.35 ± 0.08 (**Figure 5A**, cf. lane 4 to 2), indicating that Hsp47 promoted the formation of properly folded and assembled GABA_A_ receptors in the ER and increased the trafficking efficiency of the α1(A322D) variant from the ER to the Golgi. Furthermore, surface biotinylation assays demonstrated that overexpressing Hsp47 significantly increased α1(A322D) variant surface expression (**Figure 5B**). The Hsp47-enhanced α1(A322D) surface expression was also reflected in the GABA-induced *I*_max_ from 8.5 ± 4.4 pA (n = 20) to 50.3 ± 19.7 pA (n = 17) in HEK293T cells expressing α1(A322D)β2γ2 GABA_A_ receptors (**Figure 5C**). Therefore, Hsp47 promotes the functional surface expression of an epilepsy-associated GABA_A_ receptor variant.

**Figure 5.**
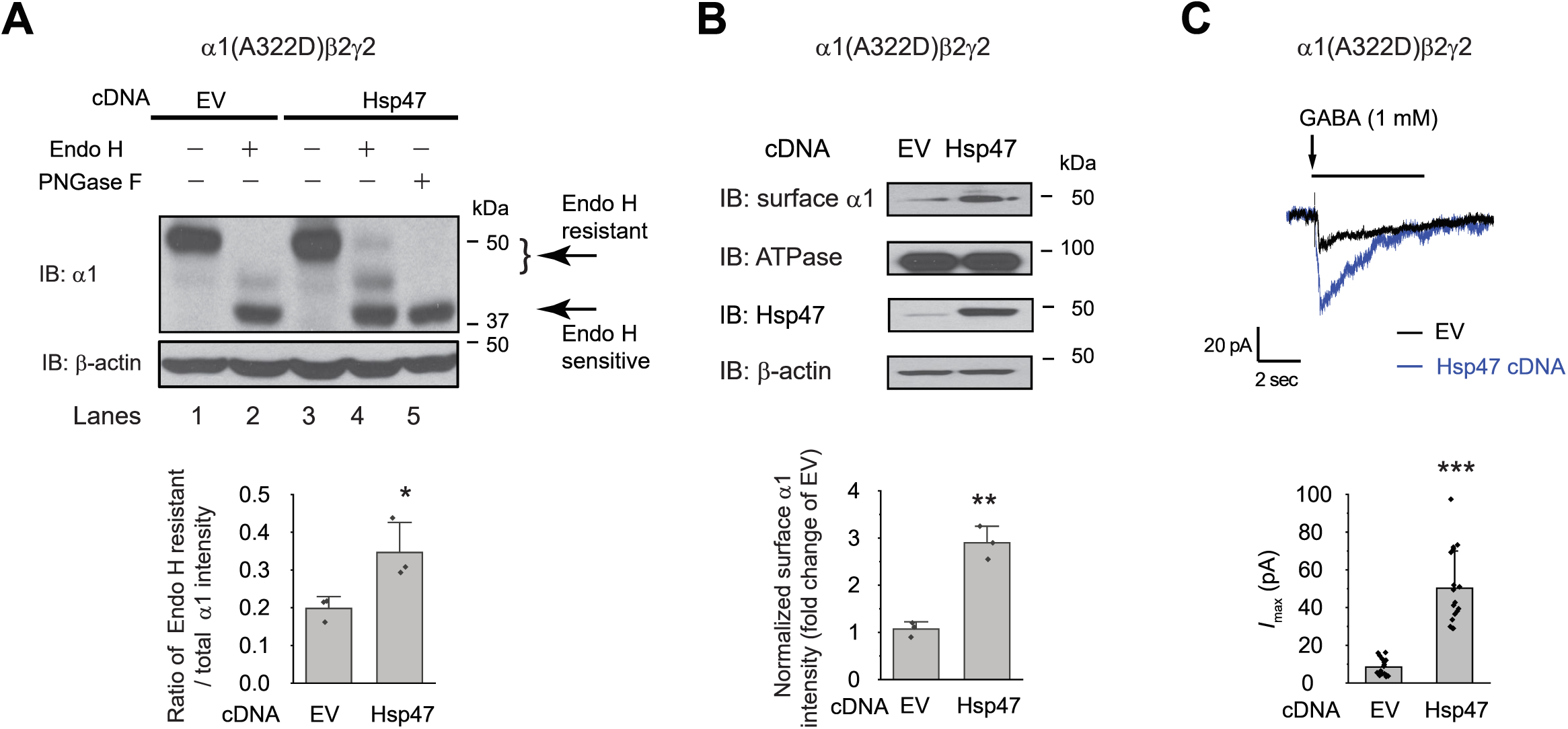
Hsp47 positively regulates the functional surface expression of epilepsy-associated GABA_A_ receptors carrying the α1(A322D) variant. (**A**) Overexpression of Hsp47 increases the endo H-resistant post-ER glycoform of the α1 subunit in HEK293T cells expressing α1(A322D)β2γ2 GABA_A_ receptors. PNGase F treatment serves as a control for unglycosylated α1 subunit (lane 5). Two endo H-resistant bands were detected for the α1 subunit, indicated by the bracket (lanes 2 and 4). Quantification of the ratio of endo H-resistant / total α1 subunit bands, as a measure of the ER-to-Golgi trafficking efficiency, is shown on the bottom (n = 3). (**B**) HEK293T cells expressing α1(A322D)β2γ2 GABA_A_ receptors were transfected with empty vector (EV) control or Hsp47 cDNA plasmids. Forty-eight hours post transfection, the surface proteins were measured using a cell surface protein biotinylation assay. The Na^+^/K^+^ ATPase serves as a loading control for biotinylated membrane proteins. Surface α1 subunit intensities were quantified using ImageJ and shown on the bottom (*n* = 3). Alternatively, cells were lysed, and the total cell lysates were subjected to SDS-PAGE and immunoblotted for Hsp47. β-actin serves as a total protein loading control. (**C**) Whole-cell patch clamping was performed to record GABA-induced currents. HEK293T cells were treated as in (**B**). The recording was carried out 48 hours post transfection. The holding potential was set at -60 mV. Representative traces were shown. Quantification of the peak currents (*I*_max_) from 17-20 cells is shown on the bottom. pA: picoampere. Each data point is reported as mean ± SD. Statistical significance was calculated using two-tailed Student’s t-Test. *, *p* < 0.05; **, *p* < 0.01; ***, *p* < 0.001.

### Hsp47 positively regulates the assembly and function of **α**4**β**2 nicotinic acetylcholine receptors (nAChRs)

The role of Hsp47 in regulating the maturation of ion channels has not been previously documented. We therefore expanded our investigation of the effect of Hsp47 to other members of the Cys-loop superfamily, to which GABA_A_ receptors belong [61]. The Cys-loop receptors, including nAChRs, are pentameric ligand-gated neuroreceptors, sharing a common structural scaffold, including a β-sheet-rich ER lumen domain [62, 63]. We chose to evaluate the effect of Hsp47 on heteropentameric α4β2 nAChRs and homopentameric α7 nAChRs since they are the major subtypes in the CNS [64]. Previously, FRET experiments were developed to evaluate the assembly of α4β2 and α7 nAChRs [58,65,66]. Here, FRET assays demonstrated that overexpressing Hsp47 significantly increased the mean FRET efficiency of (CFP-β2)(YFP-α4) nAChRs from 27.7 ± 16.4% to 40.9 ± 18.9% (**Figure 6A**), indicating that Hsp47 positively regulates the assembly of heteropentameric α4β2 receptors. In contrast, FRET experiments showed that Hsp47 overexpression did not influence the mean FRET efficiency of homopentameric α7 nAChRs (**Figure S4**), indicating that Hsp47 regulates the assembly of α4β2 and α7 nAChRs differently.

**Figure 6.**
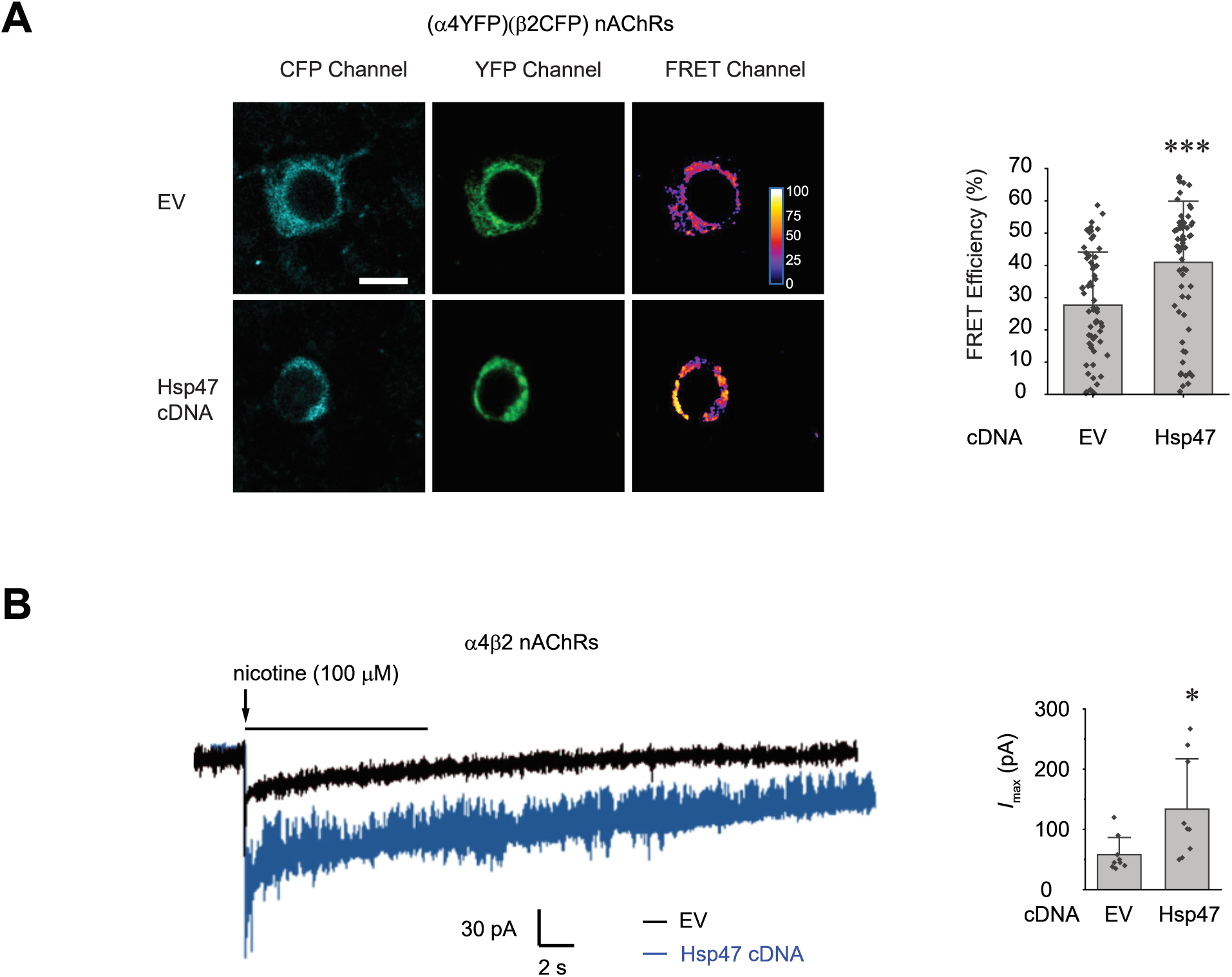
Hsp47 promotes the assembly and function of α4β2 nicotinic acetylcholine receptors (nAChRs). (**A**) Hsp47 overexpression increases FRET efficiency between CFP-tagged β2 subunit and YFP-tagged α4 subunit of nAChRs. HEK293T cells were transfected with CFP- tagged β2 subunit and YFP-tagged α4 subunit; in addition, cells were transfected with empty vector (EV) control or Hsp47 cDNA. Forty-eight hours post transfection, pixel-based FRET was used to measure the FRET efficiency between β2-CFP and α4-YFP by using a confocal microscope. Representative images were shown for the CFP channel (1st columns), YFP channel (2nd columns), and FRET efficiency (3rd columns). Scale bar = 10 μm. Quantification of the FRET efficiency from 60-70 cells was achieved using the ImageJ PixFRET plug-in, and shown on the right. (**B**) HEK293T cells were transfected with CFP-tagged β2 subunit and YFP-tagged α4 subunit of nAChRs; in addition, cells were transfected with empty vector (EV) control or Hsp47 cDNA. Forty-eight hours post transfection, whole-cell patch clamping was performed to record nicotine-induced currents. Representative traces were shown. Quantification of the peak currents (*I*_max_) from 9 cells is shown on the right. The holding potential was set at -60 mV. pA: picoampere. Each data point is reported as mean ± SD. Statistical significance was calculated using two-tailed Student’s t-Test. * *p* < 0.05; *** *p* < 0.001.

Furthermore, Hsp47 overexpression increased the nicotine-induced peak current from 57.9 ± 28.6 pA (n = 9) to 134 ± 84 pA (n = 9) in HEK293T cells expressing α4β2 nAChRs (**Figure 6B**). Therefore, Hsp47 positively regulates the assembly and thus function of α4β2 nAChRs. It appears that Hsp47 has a general role in promoting the maturation of multi-subunit neurotransmitter-gated ion channels.

## DISCUSSION

**Figure 7** illustrates our proposed mechanism for Hsp47 positively regulating the surface trafficking of GABA_A_ receptors. BiP and calnexin assist the subunit folding early in the ER lumen [49]. Hsp47 operates after BiP and binds the folded states or late folding stage of the α1 and β2 subunits in the ER lumen. Hsp47 binding links the α1 and β2 subunits and enhances their inter-subunit interactions. As a result, Hsp47 promotes the formation of the assembly intermediates and the native pentameric receptors in the ER. The composition of the assembly intermediates, such as dimers and trimers, requires future investigation. Properly assembled receptors will traffic to the Golgi and onward to the plasma membrane for function. In addition, we demonstrated that Hsp47 positively regulates the assembly and function of α4β2 nAChRs, but not the assembly of α7 nAChRs. Interestingly, it was reported that RIC-3, an ER transmembrane protein, is important for the assembly of α7, but not α4β2 nAChRs [65, 67], whereas NACHO (gene symbol: *TMEM35*), an ER transmembrane protein in the ER, can promote the assembly and function of both α4β2 and α7 nAChRs [24]. Therefore, the differential role of chaperones on the assembly of various subtypes of multi-subunit ion channels requires further investigation. Nonetheless, it appears that Hsp47 acts as a chaperone that promotes the assembly process of both GABA_A_ receptors and α4β2 nAChRs from the Cys-loop receptor family. This supplements the canonical function of molecular chaperones that serve to assist protein folding and prevent protein aggregation. The assembly of GABA_A_ receptor subunits is mediated by their N-terminal domains [51, 68]. For example, residues 86-96 within α1 subunits, especially Gln95, play an important role in their assembly with β3 subunits [51]. Here, we showed that an ER lumen localized chaperone, Hsp47, enhanced the oligomerization of hetero-pentameric GABA_A_ receptors. It would be of great interest to identify the residues that mediate the interaction between Hsp47 and GABA_A_ receptors.

**Figure 7.**
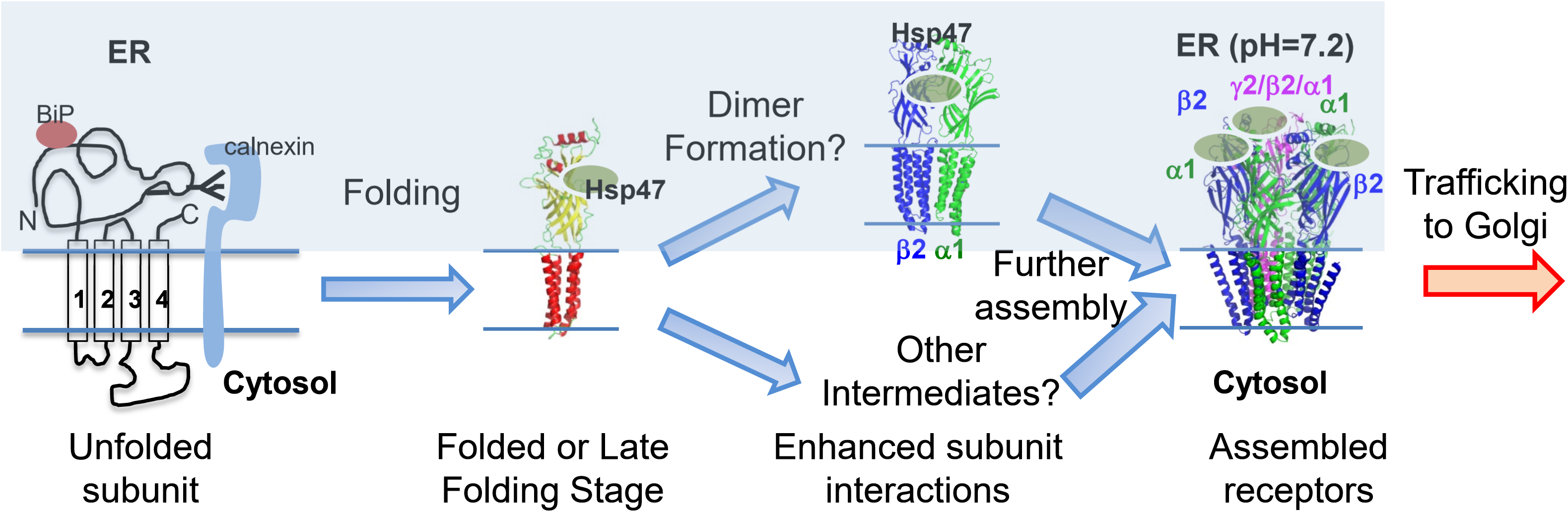
Proposed mechanism of Hsp47 in the assembly of GABA_A_ receptors. BiP and calnexin assist the subunit folding early in the ER lumen. Hsp47 operates after BiP and binds the folded states of the α1 or β subunits in the ER lumen. Hsp47 links the α1 and β subunits and promotes their inter-subunit interactions. As a result, Hsp47 promotes the formation of assembly intermediates and the native pentameric receptors in the ER. Assembled receptors will traffic to the Golgi and onward to the plasma membrane for function.

Our results expand the client protein pool and function of Hsp47. Hsp47 was identified and is currently recognized as a collagen-specific chaperone [40]. Here, we demonstrate that Hsp47 has a general effect in proteostasis maintenance of both the GABA_A_ receptor and nAChRs in the Cys-loop receptor superfamily. In addition, it was previously reported that Hsp47 physically interacts with amyloid precursor protein and regulates the secretion of Aβ-peptide [47]. Therefore, Hsp47 could have diverse functions in the CNS. Here, we show that Hsp47 binds the ER luminal domain (ERD) of GABA_A_ receptor α1 subunit with high-affinity. The GABA_A_ receptor ERD domain is rich in β-sheets containing ten β-strands [69], whereas, for its known substrate collagen, Hsp47 preferentially binds to the folded conformation of the collagen triple helices with a 2:1 stoichiometry [70–72]. It is also known that Hsp47 interacts with the ERD of IRE1α, a major transducer of the UPR, with a Kd of 73.2 ± 8.4 nM, to regulate the IRE1α oligomerization [46]; the ERD of IRE1α also adopts a β-sheet rich structure with a triangular assembly of β-sheet clusters [73]. Therefore, it appears that Hsp47 can interact with both the β-sheet-rich structure and triple helix structure. How Hsp47 adopts these structurally diverse client proteins will be the subject of future investigations. As a pentameric channel, each GABA_A_ receptor has five potential binding sites for Hsp47. However, each GABA_A_ receptor only has two agonist binding sites in the N-terminal domain for GABA. Therefore, the binding stoichiometry between Hsp47 and the pentameric GABA_A_ receptors merits future effort.

Recent advances in genetics identified mutations in GABA_A_ receptors that are associated with idiopathic epilepsies [36,38,74]. One mutation can compromise the receptor function by influencing the protein biogenesis pathways (transcription, translation, folding, assembly, trafficking, and endocytosis), ligand binding, channel gating, or their combinations. Recently, we showed that enhancing the ER folding capacity is a viable way to restore the surface expression and thus function of misfolding-prone mutations [15,49,75–78]. Numerous disease-causing mutations disrupt their folding and/or assembly, leading to reduced trafficking to the plasma membrane. These trafficking-deficient mutant subunits are retained in the ER and degraded by the ERAD pathway. Because we envisage that Hsp47 stabilizes the assembled receptor complex, overexpressing Hsp47 has the promise to enhance the forward trafficking process of such mutant receptors, consequently restoring their normal function. Indeed, we show that Hsp47 partially restored the function of the misfolding-prone α1(A322D) subunit. This strategy serves as a proof-of-principle case for promoting the multi-subunit assembly process in order to ameliorate diseases resulting from membrane protein folding/assembly deficiencies.

## MATERIALS AND METHODS

### Plasmids

The pCMV6 plasmids containing human GABA_A_ receptor α1 subunit (Uniprot #: P14867-1) (catalog #: RC205390), β2 subunit (isoform 2, Uniprot #: P47870-1) (catalog #: RC216424), γ2 subunit (isoform 2, Uniprot #: P18507-2) (catalog #: RC209260), and pCMV6 Entry Vector plasmid (pCMV6-EV) (catalog #: PS100001) were obtained from Origene. The missense mutations (A322D, C166A, or C166A/C180A in the GABA_A_ receptor α1 subunit) were constructed using a QuikChange II site-directed mutagenesis Kit (Agilent Genomics, catalog #: 200523). A FLAG tag was inserted between Leu31 and Gln32 of the α1 subunit and between Asn28 and Asp29 of the β2 subunit of GABA_A_ receptors by using QuikChange II site-directed mutagenesis. For GABA_A_ receptors, enhanced cyan fluorescent protein (CFP) was inserted between Lys364 and Asn365 in the TM3-TM4 intracellular loop of the α1 subunit, and enhanced yellow fluorescent proteins (YFP) was inserted between Lys359 and Met360 in the TM3-TM4 intracellular loop of the β2 subunit by using the GenBuilder cloning kit (GenScript, catalog #: L00701). The construction of fluorescently tagged nAChR subunits were described previously [58, 65]: CFP was inserted into the TM3-TM4 intracellular loop of the β2 subunit (Addgene, catalog #: 15106), YFP was inserted into TM3-TM4 intracellular loop of the α4 subunit (Addgene, catalog #: 15245), and cerulean (a CFP variant) or venus (a YFP variant) was inserted into TM3-TM4 intracellular loop of the α7 subunit. The human Hsp47 cDNA in pCMV6-XL5 plasmid was obtained from Origene (catalog#: SC119367). Scrambled siRNA GFP lentivector (catalog #: LV015-G) and Hsp47-set of four siRNA lentivectors (rat) (catalog #: 435050960395) were obtained from Applied Biological Materials. psPAX2 (Addgene plasmid # 12260; http://n2t.net/addgene:12260; RRID:Addgene_12260) and pMD2.G (Addgene plasmid # 12259; http://n2t.net/addgene:12259; RRID:Addgene_12259) were a gift from Didier Trono.

### Antibodies

The mouse monoclonal anti-α1 subunit antibody (clone BD24, catalog #: MAB339), mouse monoclonal anti-β2/3 subunit antibody (clone 62-3G1, catalog #: 05-474), rabbit polyclonal anti-β2 subunit antibody (catalog #: AB5561), and rabbit polyclonal anti-NeuN (catalog #: ABN78) antibody were obtained from Millipore (Burlington, MA). The rabbit polyclonal anti-α1 antibody (catalog #: PPS022) was purchased from R&D systems (Minneapolis, MN), and the goat polyclonal anti-α1 subunit antibody (A-20; catalog #: SC- 31405) was obtained from Santa Cruz Biotechnology (Dallas, TX). The mouse monoclonal anti-FLAG (catalog #: F1804) and anti-β-actin (catalog #: A1978) antibodies came from Sigma (St. Louis, MO). The mouse monoclonal anti-Hsp47 antibody (catalog #: ADI-SPA-470-F) came from Enzo Life Sciences (Farmingdale, NY). The rabbit polyclonal anti-Grp78 (BiP) antibody (catalog #: 3158-1) was obtained from Epitomics (Burlingame, CA). The rabbit polyclonal anti- Na^+^/K^+^ ATPase antibody (catalog #: AB76020) was obtained from Abcam (Waltham, MA). The mouse monoclonal anti-His tag antibody (catalog #: 2366S) was obtained from Cell Signaling Technology (Danvers, MA).

### Cell Culture and Transfection

HEK293T cells (ATCC, catalog #: CRL-3216) were maintained in Dulbecco’s Modified Eagle Medium (DMEM) (Fisher Scientific, Waltham, MA, catalog #: 10-013-CV) with 10% heat-inactivated fetal bovine serum (Fisher Scientific, catalog #: SH30396.03HI) and 1% Penicillin-Streptomycin (Fisher Scientific, catalog #: SV30010) at 37°C in 5% CO_2_. Monolayers were passaged upon reaching confluency with 0.05% trypsin protease (Fisher Scientific, catalog #: SH30236.01). Cells were grown in 6-well plates or 10 cm dishes and allowed to reach ∼70% confluency before transient transfection using TransIT-2020 (Mirus Bio, Madison, WI, catalog #: MIR 5400) according to the manufacturer’s instruction. HEK293T cells stably expressing α1β2γ2 or α1(A322D)β2γ2 GABA_A_ receptors were generated using the G418 selectin method, as described previously [15, 76]. Forty-eight hours post-transfection, cells were harvested for protein analysis.

### Lentivirus transduction in rat neurons

Lentivirus production and transduction in neurons was performed as described previously [79]. Briefly, HEK293T cells were transfected with a Hsp47-set of four siRNA lentivectors (rat) (Applied Biological Materials, catalog #: 435050960395), or scrambled siRNA GFP lentivector (Applied Biological Materials, catalog #: LV015-G) together with psPAX2 and pMD2.G plasmids using TransIT-2020. The medium was changed after 8 h incubation, and cells were incubated for additional 36-48 h. Then the medium was collected, filtered, and concentrated using Lenti-X concentrator (Takara Bio, Catalog #: 631231). The lentivirus was quantified with the qPCR lentivirus titration kit (Applied Biological Materials, catalog #: LV900), and stored at −80°C.

Sprague Dawley rat E18 hippocampus was obtained from BrainBits (Springfield, IL). Neurons were isolated and cultured following the company’s instruction. Briefly, tissues were digested with papain (2 mg/ml) (Sigma, catalog #: P4762) at 30°C for 10 min and triturated with a fire-polished sterile glass pipette for 1 min. Neurons were then plated onto poly-D-lysine (PDL) (Sigma, catalog #: P6407) and Laminin (Sigma, catalog #: L2020)-coated glass coverslips in a 24-well plate. Neurons were maintained in neuronal culture media containing Neurobasal (ThermoFisher, catalog #: 21103049), 2% B27 (ThermoFisher, catalog #: 17504044), 0.5 mM GlutaMax (ThermoFisher, catalog #: 35050061), and 1% penicillin-streptomycin (Fisher Scientific, catalog #: SV30010) at 37°C in 5% CO_2_. Neurons were subjected to transduction with lentivirus at days *in vitro* (DIV) 10, and immunofluorescence staining and electrophysiology were performed at DIV 12.

### Mouse brain homogenization

C57BL/6J mice (Jackson Laboratory) at 8-10 weeks were sacrificed and the cortex was isolated and homogenized in the homogenization buffer (25 mM Tris, pH 7.6, 150 mM NaCl, 1 mM EDTA, and 2% Triton X-100) supplemented with the Roche complete protease inhibitor cocktail (Roche; catalog #: 4693159001). The homogenates were centrifuged at 800 × g for 10 min at 4 °C, and the supernatants were collected. The pellet was re-homogenized in additional homogenization buffer and centrifuged at 800 × g for 10 min at 4 °C. The supernatants were combined and rotated for 2 h at 4 °C, and then centrifuged at 15,000 × g for 30 min at 4 °C. The resulting supernatant was collected as mouse brain homogenate. Protein concentration was determined by a MicroBCA assay (Pierce, catalog #: 23235). This animal study was approved by the Institutional Animal Care and Use Committees (IACUC) at Case Western Reserve University and was carried out in agreement with the recommendation of the American Veterinary Medical Association Panel on Euthanasia.

### Western Blot Analysis

Cells were harvested and lysed with lysis buffer (50 mM Tris, pH 7.5, 150 mM NaCl, and 1% Triton X-100) or RIPA buffer (50 mM Tris, pH 7.4, 150 mM NaCl, 5 mM EDTA, pH 8.0, 2% NP-40, 0.5% sodium deoxycholate, and 0.1% SDS) supplemented with Roche complete protease inhibitor cocktail. Lysates were cleared by centrifugation (20,000 × *g*, 10 min, 4 °C).

Protein concentration was determined by MicroBCA assay. Aliquots of cell lysates were separated in an 8% SDS-PAGE gel, and Western blot analysis was performed using the appropriate antibodies. Band intensity was quantified using ImageJ software from the National Institute of Health [80]. For non-reducing protein gels, cell lysates were loaded in the Laemmli sample buffer (Biorad, Hercules, CA, catalog #: 1610737); for reducing protein gels, cell lysates were loaded in the Laemmli sample buffer (Biorad, catalog #: 1610737) supplemented with 100 mM dithiothreitol (DTT) to reduce the disulfide bonds. Endoglycosidase H (endo H) (New England Biolabs, catalog #: P0703L) enzyme digestion or Peptide-N-Glycosidase F (PNGase F) (New England Biolabs, Ipswich, MA, catalog #: P0704L) enzyme digestion was performed according to manufacturer’s instruction and the published procedure [49].

### Immunoprecipitation

Cell lysates (500 µg) or mouse brain homogenates (1 mg) were pre-cleared with 30 µL of protein A/G plus-agarose beads (Santa Cruz Biotechnology, catalog #: SC-2003) and 1.0 µg of normal mouse IgG (Santa Cruz Biotechnology, catalog #: SC-2025) for 1 hour at 4°C to remove nonspecific binding proteins. The pre-cleared samples were incubated with 2.0 µg of mouse anti-

α1 antibody, mouse anti-Hsp47 antibody, or normal mouse IgG as a negative control for 1 hour at 4°C and then with 30 µL of protein A/G plus agarose beads overnight at 4°C. Afterward, the beads were collected by centrifugation at 8000 ×g for 30 s, and washed three times with lysis buffer. The complex was eluted by incubation with 30 µL of Laemmli sample buffer loading buffer in the presence of 100 mM DTT. The immunopurified eluents were separated in an 8% SDS-PAGE gel, and Western blot analysis was performed using appropriate antibodies.

### *In vitro* protein binding assay

1 μg of GST epitope tag protein (GST) (Novus Biologicals, Centennial, CO, catalog #: NBC1-18537), GST-tagged human GABA_A_ receptor α1 subunit protein (GST-α1) (Abnova, Walnut, CA, catalog #: H00002554-P01), or GST-tagged human GABA_A_ receptor β2 subunit protein (GST-β2) (Abnova, catalog #: H00002561-P01) was mixed with 4 μg of recombinant His-tagged human Hsp47 protein (Novus, catalog #: NBC1-22576) in 500 µL of lysis buffer (50 mM Tris, pH 7.5, 150 mM NaCl, and 1% Triton X-100). The protein complex was isolated by immunoprecipitation using a mouse anti-His antibody (Cell Signaling, catalog #: 2366S) followed by SDS-PAGE and Western blot analysis with a rabbit anti-GABA_A_ receptor α1 subunit antibody (R&D Systems, catalog #: PS022), a mouse anti-GABA_A_ receptor β2/β3 subunit antibody (Millipore, catalog #: 05-474), or a mouse anti-His antibody.

### MicroScale Thermophoresis (MST)

MST experiments were carried out to measure the binding affinity between the ER luminal domain of human GABA_A_ receptor α1 subunits (α1-ERD) and ER luminal chaperones, Hsp47 and BiP, using a Monolith NT.115 instrument (NanoTemper Technologies Inc, South San Francisco, CA). Monolith His-Tag Labeling Kit RED-tris-NTA 2^nd^ Generation (NanoTemper Technologies, catalog #: MO-L018) was used to label recombinant His-α1-ERD (MyBioSource, San Diego. CA, catalog #: MBS948971). 100 μL of 200 nM RED-tris-NTA dye in PBST buffer (137 mM NaCl, 2.7 mM KCl, 10 mM Na_2_HPO_4_, 1.8 mM KH_2_PO_4_, pH 7.4, 0.05% Tween-20) was mixed with 100 μL of 200 nM His-α1-ERD in PBST, and the reaction mixture was incubated for 30 min at room temperature in the dark. The serial dilutions of the ligand proteins (0.2 nM to 10 μM) in PBST, including recombinant human Hsp47 (Abcam, catalog #: ab86918) and human BiP (Abcam, catalog #: ab78432), were prepared in Maxymum Recovery PCR tubes (Axygen, Union City, CA, catalog #: PCR-02-L-C) with a final volume of 5 μL in each tube. Then 5 μL of 100 nM RED labeled His-α1-ERD was added to each PCR tube containing 5 μL of the ligand proteins. The samples were loaded to the capillaries (NanoTemper Technologies, catalog #: MO-K025) and measured using a Monolith NT.115 instrument with the settings of 40% LED/excitation and 40% MST power. The data were collected and analyzed using the Monolith software for the calculation of the dissociation constant (Kd).

### Biotinylation of Cell Surface Proteins

Cells were plated in 6-cm dishes for surface biotinylation experiments according to the published procedure [49]. Briefly, intact cells were washed twice with ice-cold Dulbecco’s phosphate buffered saline (DPBS) (Fisher Scientific, catalog #: SH3002803). To label surface membrane proteins, cells were incubated with the membrane-impermeable biotinylation reagent Sulfo-NHS SS-Biotin (0.5 mg ⁄ mL) (Pierce, catalog #: 21331) in DPBS containing 0.1 mM CaCl_2_ and 1 mM MgCl_2_ (DPBS+CM) for 30 min at 4 °C. To quench the reaction, cells were incubated with 10 mM glycine in ice-cold DPBS+CM twice for 5 min at 4 °C. Sulfhydryl groups were blocked by incubating the cells with 5 nM N-ethylmaleimide (NEM) (Pierce, catalog #: 23030) in DPBS for 15 min at room temperature. Cells were then solubilized for 1 h at 4 °C in solubilization buffer (50 mM Tris–HCl, 150 mM NaCl, 5 mM EDTA, pH 7.5, 1% Triton X-100) supplemented with Roche complete protease inhibitor cocktail and 5 mM NEM. The samples were centrifuged at 20,000 × g for 10 min at 4 °C to pellet cellular debris. The supernatant contained the biotinylated surface proteins. The concentration of the supernatant was measured using microBCA assay. Biotinylated surface proteins were affinity-purified from the above supernatant by incubating for 1 h at 4 °C with 100 μL of immobilized neutravidin-conjugated agarose bead slurry (Pierce, catalog #: 29200). The samples were then centrifuged at 20,000 ×g for 10 min at 4 °C. The beads were washed six times with solubilization buffer. Surface proteins were eluted from beads by boiling for 5 min with 200 μL of LSB ⁄ Urea buffer (2x Laemmli sample buffer (LSB) with 100 mM DTT and 6 M urea; pH 6.8) for SDS-PAGE and Western blotting analysis.

### Immunofluorescence Staining and Confocal Microscopy

Neuron staining and confocal immunofluorescence microscopy analysis were performed as described previously [49, 79]. Briefly, to label cell surface proteins, primary neurons on coverslips were fixed with 2% paraformaldehyde in DPBS for 10 min. We then blocked with 10% goat serum (ThermoFisher, catalog #: 16210064) in DPBS for 0.5 h, and without detergent permeabilization, incubated with 100 μL of appropriate primary antibodies against the GABA_A_ receptor α1 subunit (Synaptic Systems, Goettingen, Germany, catalog #: 224203) (1:250 dilution) or β2/3 subunits (Millipore, catalog #: 05-474) (1:250 dilution), diluted in 2% goat serum in DPBS, at room temperature for 1 h. Then the neurons were incubated with Alexa 594- conjugated goat anti-rabbit antibody (ThermoFisher, catalog #: A11037), or Alexa 594- conjugated goat anti-mouse antibody (ThermoFisher, catalog #: A11032) (1:500 dilution) diluted in 2% goat serum in DPBS for 1 h. Afterward, cells were permeabilized with saponin (0.2%) for 5 min and incubated with DAPI (1 μg/mL) (ThermoFisher, catalog #: D1306) for 3 min to stain the nucleus. To label intracellular proteins, neurons were fixed with 4% paraformaldehyde in DPBS for 15 min, permeabilized with saponin (0.2%) in DPBS for 15 min, and blocked with 10% goat serum in DPBS for 0.5 h at room temperature. Then neurons were labeled with appropriate primary antibodies against Hsp47 (Enzo Life Sciences, catalog #: ADI-SPA-470-F) (1:100 dilution) or NeuN, a neuron nuclei marker (Millipore, catalog #: ABN78) (1:500 dilution), diluted in 2% goat serum in DPBS, for 1 h. The neurons were then incubated with Alexa 594-conjugated goat anti-mouse antibody (ThermoFisher, catalog #: A11032) (1:500 dilution), or Alexa 350-conjugated goat anti-rabbit antibody (ThermoFisher, catalog #: A21068) (1:500 dilution), diluted in 2% goat serum in DPBS, for 1 h. The coverslips were then mounted using fluoromount-G (VWR, catalog #: 100502-406) and sealed. An Olympus IX-81 Fluoview FV1000 confocal laser scanning system was used for imaging with a 60× objective by using FV10-ASW software. The images were analyzed using ImageJ software [80].

### Pixel-based sensitized acceptor emission FRET microscopy

Pixel-by-pixel based sensitized acceptor FRET microscopy was performed as described previously[65, 81]. For FRET experiments on GABA_A_ receptors, (1) for FRET pair samples, HEK293T cells on coverslips were transfected with α1-CFP (donor) (0.5 μg), β2-YFP (acceptor) (0.5 μg), and γ2 (0.5 μg) subunits; (2) for the donor-only samples, to determine the spectral bleed-through (SBT) parameter for the donor, HEK293T cells were transfected with α1-CFP (donor) (0.5 μg), β2 (0.5 μg), and γ2 (0.5 μg) subunits; (3) for the acceptor-only samples, to determine the SBT parameter for the acceptor, HEK293T cells were transfected with α1 (0.5 μg), β2-YFP (acceptor) (0.5 μg), and γ2 (0.5 μg) subunits. For FRET experiments on nAChRs, HEK293T cells were transfected with β2-CFP (donor) (0.7 μg) and α4-YFP (acceptor) (0.7 μg) or α7-cerulean (donor) (0.7 μg) and α7-venus (acceptor) (0.7 μg); in addition, the donor-only samples or the acceptor-only samples were prepared to determine the SBT parameters for the donor or the acceptor, respectively. Whole-cell patch-clamp recordings in HEK293T cells showed that fluorescently tagged ion channels have similar peak current amplitudes and dose-response curves to the agonists as compared to untagged ion channels (**Figure S3**) [65]. The coverslips were then mounted using fluoromount-G and sealed. An Olympus Fluoview FV1000 confocal laser scanning system was used for imaging with a 60× 1.35 numerical aperture oil objective by using Olympus FV10-ASW software.

For the FRET pair samples, donor images were acquired at an excitation wavelength of 433 nm and an emission wavelength of 478 nm, FRET images at 433 nm excitation and 528 nm emission wavelengths, and acceptor images at 514 nm excitation and 528 nm emission wavelengths. For the donor-only samples, donor images were acquired at 433 nm excitation and 478 nm emission wavelengths, and FRET images at 433 nm excitation and 528 nm emission wavelengths. For the acceptor-only samples, FRET images were acquired at an excitation of 433 nm and an emission of 528 nm, and acceptor images at an excitation of 514 nm and an emission of 528 nm. Image analysis of FRET efficiencies was performed using the PixFRET plugin of the ImageJ software [82]. The bleed-through was determined for the donor and the acceptor. With the background and bleed-through correction, the net FRET (nFRET) was calculated according to equation (1).

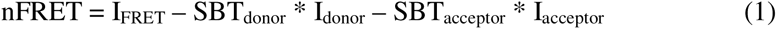

FRET efficiencies from sensitized emission experiments were calculated according to equation (2).

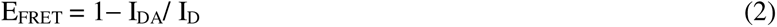

EFRET represents FRET efficiency, IDA represents the emission intensity of the donor in the presence of the acceptor, and ID represents the emission intensity of the donor alone. Since ID can be estimated by adding the nFRET signal amplitude to the amplitude of IDA [83], FRET efficiency was calculated according to equation (3).

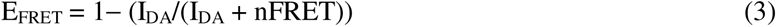

### Whole-Cell Patch-Clamp Electrophysiology

Whole-cell patch-clamp recording was performed at room temperature, as described previously for GABA_A_ receptors [77] or nAChRs [84]. Briefly, the glass electrodes have a tip resistance of 3–5 MΩ when filled with intracellular solution. For GABA_A_ receptor recording, the intracellular solution was composed of (in mM): 153 KCl, 1 MgCl_2_, 5 EGTA, 10 HEPES, and 5 Mg-ATP (adjusted to pH 7.3 with KOH); the extracellular solution was composed of (in mM): 142 NaCl, 8 KCl, 6 MgCl_2_, 1 CaCl_2_, 10 glucose, 10 HEPES, and 120 nM fenvalerate (adjusted to pH 7.4 with NaOH). For nAChR recording, the intracellular solution was composed of (in mM): 135 K-gluconate, 5 KCl, 5 EGTA, 0.5 CaCl_2_, 10 HEPES, 2 Mg-ATP, and 0.1 GTP (adjusted to pH 7.2 with Tris-base); the extracellular solution was composed of (in mM): 140 NaCl, 5 KCl, 2 CaCl_2_, 1 MgCl_2_, 10 HEPES, and 10 glucose (adjusted to pH 7.3 with Tris-base). For GABA_A_ receptor recordings, coverslips containing cells were placed in a RC-25 recording chamber (Warner Instruments) on the stage of an Olympus IX-71 inverted fluorescence microscope and perfused with extracellular solution. Fast chemical application was accomplished with a pressure-controlled perfusion system (Warner Instruments) positioned within 50 µm of the cell utilizing a Quartz MicroManifold with 100-µm inner diameter inlet tubes (ALA Scientific). The whole-cell currents were recorded at a holding potential of −60 mV in voltage-clamp mode using an Axopatch 200B amplifier (Molecular Devices, San Jose, CA). The signals were acquired at 10 kHz and filtered at 2 kHz using pClamp10 software (Molecular Devices). For nAChR recordings, cells were visualized with an upright microscope (Axio Examiner A1, Zeiss) equipped with an Axiocam 702 mono camera. Whole-cell currents were recorded at a holding potential of -60 mV in voltage clamp mode using an Integrated Patch-Clamp Amplifier (Sutter). Signals were detected at 10 kHz and filtered at 2 kHz using SutterPatch acquisition software.

## QUANTIFICATION AND STATISTICAL ANALYSIS

All data are presented as mean ± SD. If two groups were compared, statistical significance was calculated using an unpaired Student’s t-test; if more than two groups were compared, we used ANOVA followed by post hoc Tukey. A p< 0.05 was considered statistically significant. *, p< 0.05; **, p< 0.01; ***, p< 0.001.

## SUPPLEMENTAL INFORMATION

Supplemental Information includes four figures.

## ACKNOWLEDGMENTS

This work was supported by the National Institutes of Health (R01NS105789 and R01NS117176 to TM). We thank Dr. Matthias Buck (Case Western Reserve University, Cleveland, Ohio) for the help from his group on the MST experiments.

## AUTHOR CONTRIBUTIONS

Conceptualization, YW and TM; Data curation: YW, XD, DH, RN, BH, and FM; Formal analysis: YW, XD, DH, RN, BH, FM, and TM; Funding acquisition: TM; Supervision: TM; Writing – original draft: YW, XD, and TM; Writing – review & editing: YW, XD, RN, BH, FM, and TM.

