## Supplemental Figures for "Hsp47 Promotes Biogenesis of Multi-subunit Neuroreceptors in the Endoplasmic Reticulum"

### SUPPLEMENTAL INFORMATION

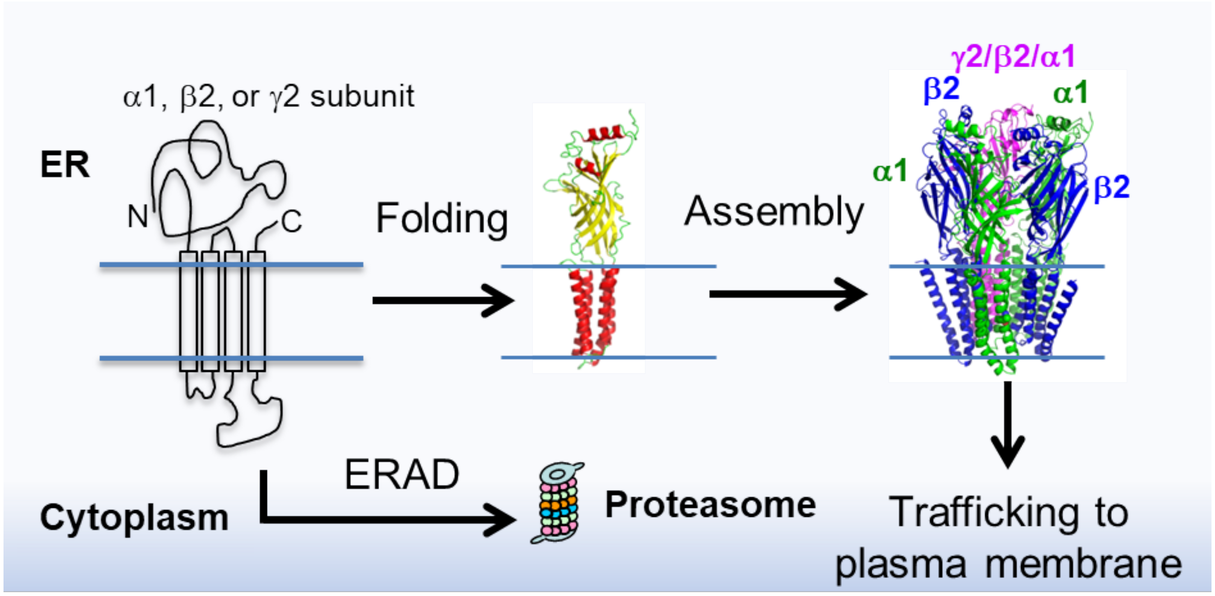


**Figure S1.** The $GABA_A$ receptor biogenesis pathway. Individual $GABA_A$ receptor subunits fold in the endoplasmic reticulum (ER). Properly folded subunits assemble into a heteropentamer in the ER for subsequent trafficking to the plasma membrane. Unassembled and misfolded subunits are degraded by the ER-associated degradation (ERAD) pathway. The $GABA_A$ receptor cartoons are built from the crystal structure of a human β3 subunit (PDB: 4COF). The large intracellular domain (ICD) is missing in the crystal structure.

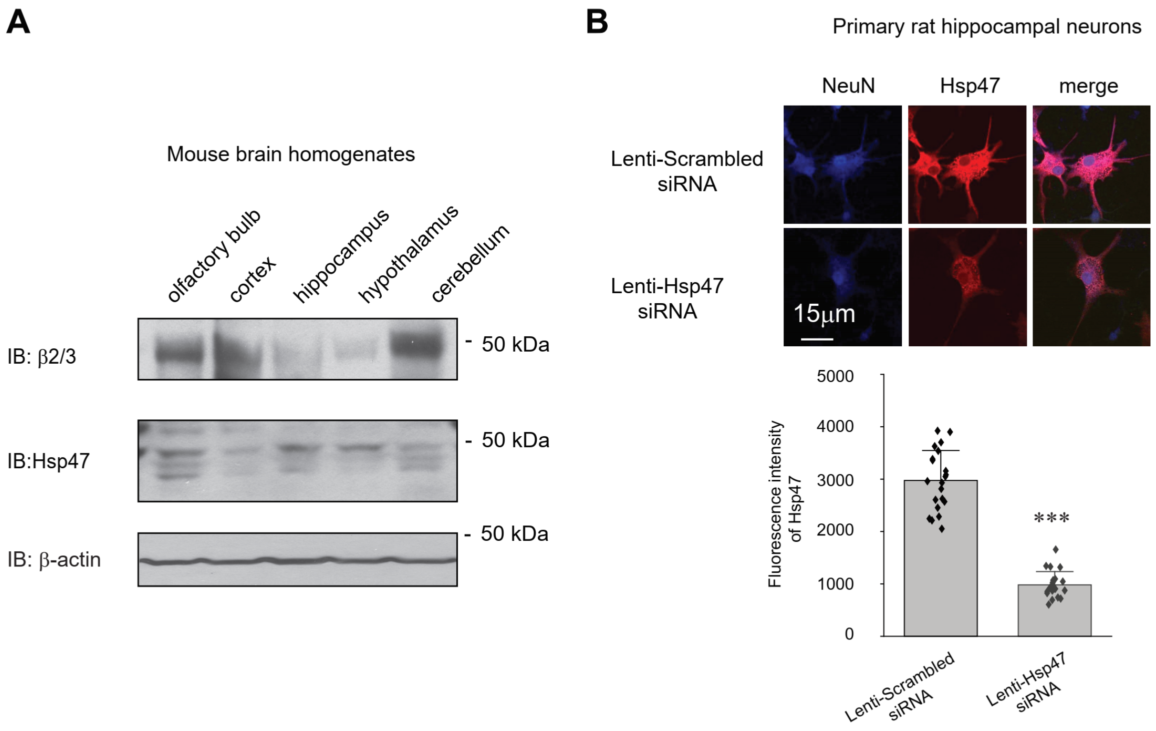


**Figure S2.** Hsp47 expression in the central nervous system. (**A**) Hsp47 and $GABA_A$ receptor β2/β3 subunit protein expression in various mouse brain regions according to SDS-PAGE and Western blot analysis. Three replicate experiments were performed from tissue isolated from three different mice for each brain region. (**B**) Hsp47 knockdown in cultured rat hippocampal neurons. Cultured neurons were subjected to transduction with Hsp47 siRNA lentivirus or scrambled siRNA lentivirus at days *in vitro* (DIV) 10. Forty-eight hours post-transduction, neurons were fixed, permeabilized, and stained using anti-Hsp47 or anti-NeuN (a marker of the neuron nuclei) antibodies. Neurons were visualized using a confocal microscope. Representative images are shown for each condition. Scale bar = 15 μm. in the bottom panel, we display the quantification of the Hsp47 staining fluorescence intensity after background correction. The analysis was performed on at least 20 cells accumulated from a minimum of three individual coverslips from either the Hsp47 siRNA lentivirus or scrambled siRNA lentivirus conditions.

Each data point is reported as mean ± SD. Statistical significance was calculated using an unpaired two-tailed Student's t-Test. *** $p < 0.001$.

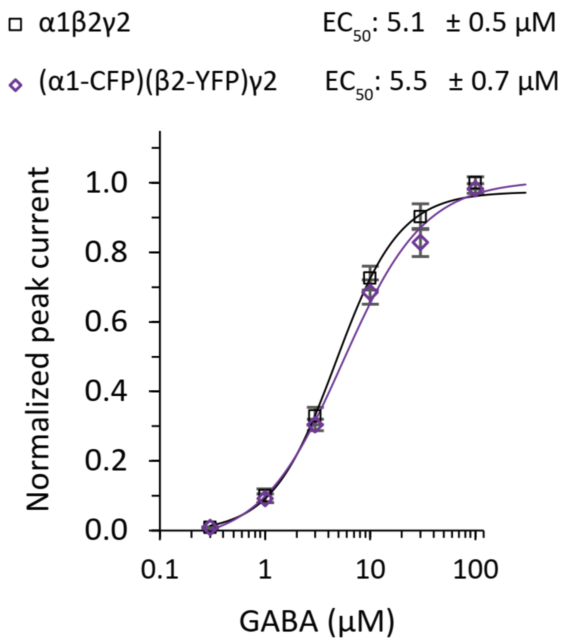


**Figure S3.** Dose-response curves of $GABA_A$ receptors. HEK293T cells were transfected with α1, β2, and γ2 subunits of $GABA_A$ receptors, or α1-CFP, β2-YFP, and γ2 subunits. 48 hours post-transfection, whole-cell patch-clamping electrophysiological recordings were carried out to calculate $EC_{50}$ values for GABA. n = 5 to 8. The holding potential was set at -60 mV. Each data point is reported as mean ± SD.

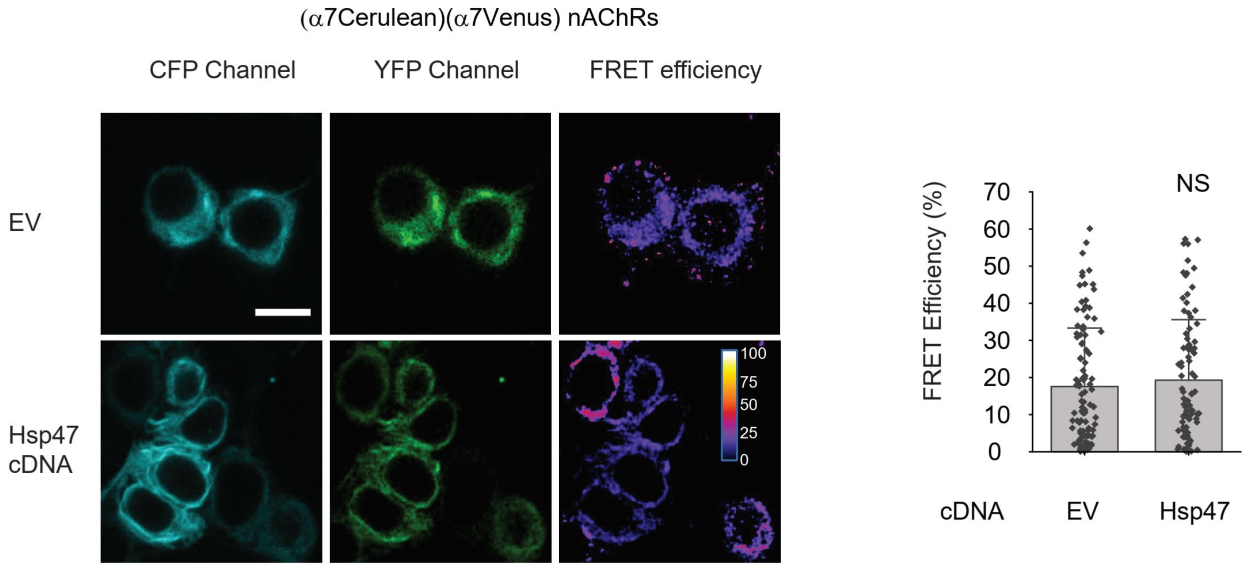


**Figure S4.** Hsp47 overexpression does not change the FRET efficiency between Cerulean (an improved CFP variant)-tagged α7 subunit and Venus (an improved YFP variant)-tagged α7 subunit of nAChRs. HEK293T cells were transfected with Cerulean-tagged α7 subunit and Venus-tagged α7 subunit; in addition, cells were transfected with empty vector (EV) control or Hsp47 cDNA. Forty-eight hours post transfection, pixel-based FRET was used to measure the FRET efficiency between α7-Cerulean and α7-Venus by using a confocal microscope. Representative images were shown for the CFP channel (1st columns), YFP channel (2nd columns), and FRET efficiency (3rd columns). Scale bar = 10 μm. Quantification of the FRET efficiency from 90-105 cells from at least three transfections was achieved using the ImageJ PixFRET plug-in, and shown on the right. Each data point is reported as mean ± SD. Statistical significance was calculated using two-tailed Student's t-Test. NS, not significant.
